# TREM2 alters the phagocytic, apoptotic and inflammatory response to Aβ_42_ in HMC3 cells

**DOI:** 10.1101/2020.10.08.329938

**Authors:** Rumana Akhter, Yvonne Shao, Shane Formica, Maria Khrestian, Lynn M. Bekris

**Affiliations:** Genomic Medicine Institute, Lerner Research Institute, Cleveland Clinic Foundation, Cleveland, OH, USA

**Keywords:** Aβ, TREM2, Microglia, Phagocytosis, Apoptosis, Inflammatory factors

## Abstract

Alzheimer’s disease (AD) is characterized by the accumulation in the brain of extracellular amyloid β (Aβ) plaques as well as intraneuronal inclusions (neurofibrillary tangles) consisting of total tau and phosphorylated tau. Also present are dystrophic neurites, loss of synapses, neuronal death, and gliosis. AD genetic studies have highlighted the importance of inflammation in this disease by identifying several risk associated immune response genes, including TREM2. TREM2 has been strongly implicated in basic microglia function including, phagocytosis, apoptosis, and the inflammatory response to Aβ in mouse brain and primary cells. These studies show that microglia are key players in the response to Aβ and in the accumulation of AD pathology. However, details are still missing about which apoptotic or inflammatory factors rely on TREM2 in their response to Aβ, especially in human cell lines. Given these previous findings our hypothesis is that TREM2 influences the response to Aβ toxicity by enhancing phagocytosis and inhibiting both the BCL-2 family of apoptotic proteins and pro-inflammatory cytokines. Aβ_42_ treatment of the human microglial cell line, HMC3 cells, was performed and TREM2 was overexpressed or silenced and the phagocytosis, apoptosis and inflammatory response were evaluated. Results indicate that a robust phagocytic response to Aβ after 24 hours requires TREM2 in HMC3 cells. Also, TREM2 inhibits Aβ induced apoptosis by activating the Mcl-1/Bim complex. TREM2 is involved in activation of IP-10, MIP-1a, and IL-8, while it inhibits FGF-2, VEGF and GRO. Taken together, TREM2 plays a role in enhancing the microglial functional response to Aβ toxicity in HMC3 cells. This novel information suggests that therapeutic strategies that seek to activate TREM2 may not only enhance phagocytosis, but it may also inhibit beneficial inflammatory factors, emphasizing the need to define TREM2-related inflammatory activity in not only mouse models of AD, but also in human AD.

## 1. Introduction

Alzheimer’s disease (AD) is the leading cause of dementia worldwide. (Alzheimer’s 2016) The histopathological hallmarks of AD are extracellular amyloid plaques composed predominantly of the amyloid-β 1-42 (Aβ_42_) peptide and neurofibrillary tangles (NFTs) within neurons consisting of abnormally aggregated, hyperphosphorylated tau protein. (Castellani et al. 2010, Serrano-Pozo et al. 2011) Microglia are the resident innate immune cells in the central nervous system (CNS) that survey the surrounding environment. Genome-wide association studies (GWAS) have identified the loss of function genetic variants of triggering receptor expressed on myeloid cell 2 (TREM2), immunoreceptor primarily present on microglia, are associated with Alzheimer’s disease risk, in which chronic inflammatory responses can occur. (Guerreiro et al. 2013, Jonsson et al. 2013, Abduljaleel et al. 2014, Gallardo and Holtzman 2019)

Microglia survey the brain for abnormalities and are rapidly activated to phagocytose cellular debris and are thus key players in maintaining brain homeostasis (Priller and Prinz 2019, Afridi et al. 2020). Microglial cells come in close proximity of the senile plaques in AD and undergoes morphological change from resting ramified appearance to an activated amoeboid phenotype (Mandrekar-Colucci and Landreth 2010). The microglial phenotypic plasticity and the impact of microglial phagocytosis play important role in disease progression (Lull and Block 2010). Mouse models and mouse microglia studies have revealed that TREM2 is involved in a variety of physiological processes, such as phagocytosis of Aβ and the pro-inflammatory response. (Gao et al. 2013, Sieber et al. 2013, Kawabori et al. 2015) TREM2 suppresses the pro-inflammatory response (Takahashi et al. 2005, Takahashi et al. 2007) and TREM2 deletion abolishes Aβ-induced microglial proliferation and apoptosis. (Zhao et al. 2018)

Apoptosis is an essential physiological process that plays a critical role tissue homeostasis. Atleast two major apoptotic pathways have been described in mammalian cells: the intrinsic pathway via Caspase-9 activation, in which death arises from mitochondrial dysfunction, and the extrinsic pathway via Caspase-8 activation, in which death is initiated from the activation of cell surface receptors. Bcl-2 family members play an important role in the intrinsic apoptotic pathway in several AD models (Paradis et al. 1996, Akhter et al. 2015, Akhter et al. 2018). The Bcl-2 family constitutes pro-survival (*e.g.* Bcl-2 and BclxL), multidomain proapoptotic (*e.g.* BAX and BAK), and BH3-only proapoptotic (*e.g.* Bim, Puma, Bid) proteins (Youle and Strasser 2008, Akhter et al. 2014). Interactions of Bcl-2 family members function as a ‘life/death switch’ that integrates diverse inter- and intracellular cues to determine whether or not the stress apoptosis pathway should be activated. The Bcl-2 family has been involved in the microglia response to Aβ. (Shang et al. 2012) The Bcl family of proteins have been implicated in the Bcl family Akt signaling response to amyloid. (Clementi et al. 2006, Yin et al. 2011, Zhu et al. 2015) However, even though TREM2-related Akt signaling has been described, (He et al. 2019, Chen et al. 2020) to our knowledge a link between TREM2 specific signaling and Bcl-2 family mediated apoptosis has not been studied. In addition, TREM2 inhibits the inflammatory response to LPS in mouse microglia by suppressing the PI3K/NF-κB signaling, (Li et al. 2019) but less is known about the response to Aβ in human microglia.

TREM2 inhibits the inflammatory response to LPS in mouse microglia by suppressing the PI3K/NF-κB signaling, (Li et al. 2019) but less is known about the response to Aβ in human microglia. Increased expression of TREM2 upon PRRSV infection in vitro. TREM2 silencing restrained the replication of PRRSV, whereas TREM2 overexpression facilitated viral replication. The cytoplasmic tail domain of TREM2 interacted with PRRSV Nsp2 to promote infection. TREM2 downregulation led to early activation of PI3K/NF-κB signaling, thus reinforcing the expression of proinflammatory cytokines and type I interferons. (Zhu et al. 2020) TREM2 overexpression downregulation the expression of the TLR family (TLR2, TLR4 and TLR6) in BV-2 cells. Moreover, LPS, as an agonist of the TLR family, up-regulated the expression of inflammatory cytokines (IL-1β, TNF-α, and IL-6) in BV-2 cells overexpressing TREM2. (Long et al. 2019) The levels of TREM2, IL-6, and TNF-α significantly increased in peripheral blood of spinal cord injury patients and in LPS-stimulated mouse primary microglia cells. Down-regulation of TREM2 significantly reduced the expression of p-NF-κB and the production of IL-6 and TNF-α. In addition, PDTC (a NF-κB inhibitor) treatment inhibited TREM2 overexpression induced-production of IL-6 and TNF-α. (Wang et al. 2019) LPS stimulation of BV2 microglia significantly increased the production of TNF-α, IL-1β, and the activation of AKT and NF-kB. In contrast, it decreased the levels of IL-10 and TGF-β1. However, these pro-inflammatory effects were significantly attenuated by TREM2 overexpression, while enhanced by TREM2 silencing. (Li et al. 2019) The cleaved soluble TREM2 isoform has been associated with alterations in the immune response in Down syndrome (Weber et al. 2020), after exercise (Jensen et al. 2019), and AD. (Bekris et al. 2018, Brosseron et al. 2018, Rauchmann et al. 2020) Interestingly, CSF sTREM2 was positively associated with the pro-inflammatory proteins and anti-inflammatory proteins. (Rauchmann et al. 2020) Indeed, it has been suggested that sTREM2 cerebrospinal fluid levels are a potential biomarker for microglia activity in early-stage Alzheimer’s disease and associated with neuronal injury markers.(Suárez-Calvet et al. 2016) However, which inflammatory factors are influenced by TREM2 after Aβ treatment has not been fully elucidated.

Therefore, the aim of this study is to identify key factors influenced by TREM2 in the response to Aβ_42_ treatment. Given previous evidence the hypothesis is that TREM2 enhances the response to Aβ toxicity by enhancing phagocytosis and inhibiting both the Bcl family of apoptotic proteins and pro-inflammatory cytokines. In this investigation three critical microglia activities, phagocytosis, the production of intrinsic apoptotic proteins and the production of inflammatory factors were evaluated to understand whether TREM2 alters the response to Aβ toxicity in a human microglia cell line. Indeed, the results suggest that phagocytic response to Aβ requires TREM2.

## 2. Materials and Methods

### Cell Culture

Immortalized human cell lines were obtained from American Type Culture Collection (ATCC®) included a hepatocyte cell line; HepG2, two neuronal cell lines; IMR-32, CHP-212, three glial cell lines; U87, U118, HMC3 and two monocyte cell lines; THP-1, U937. HepG2, IMR-32, U87, U118, U138 cell lines were cultured in DMEM medium, HMC3 with EMEM, THP-1 with RPMI and U937 with RPMI-1640. All cells line culture conditions also included 10% FBS and 1% penicillin/streptomycin at 37 °C in a humidified atmosphere containing 95% air and 5% CO2, and the medium was changed every two days. The cells were split with 0.05% trypsin when they reached 80% confluence and sub-cultured for further passages. HMC3 (ATCC®: CRL-3304™)(Janabi et al. 1995, Li et al. 2009, Jadhav et al. 2014, Nakagawa and Chiba 2015, Dello Russo et al. 2018)

### Quantitative RT-PCR and Immunoblot Analysis

RNA and protein were extracted from cell lines using the Qiagen AllPrep DNA/RNA Mini Kit (Qiagen, Valencia, CA, USA). Total RNA was also DNase treated using the TURBO DNA-free Kit (Applied Biosystems, Austin, TX, USA) to further deplete DNA content. Protein was extracted using a modification of the Qiagen AllPrep DNA/RNA Mini Kit (Qiagen) according to manufacturer instructions. Total cell line extracts (36 μg protein) from the different treatments were separated by 12.5% SDS-PAGE and electroblotted onto supported nitrocellulose. Each blot was blocked for 1 h in Tris-buffered saline/Tween-20 (10 mM Tris base, 150 mM NaCl, 0.05% Tween-20) containing 5% fat-free milk, and then incubated with a primary TREM2 antibody (TREM2 Ab MAB1828: R&D Systems) overnight at 4°C. Washing of membranes three times (10 min each) with Tris-buffered saline/Tween-20 was followed by incubation for 1 h at room temperature with the appropriate secondary antibody (Anti-mouse IgG horseradish peroxidase cat# NA931V from GE Healthcare, Uk). Western blots were developed using the ECL Prime western blotting detection Reagent (RPN 2232: GE Healthcare). was reprobed with anti-actin (Actin A4700 mouse monoclonal: Sigma) to verify that proteins were uniformly loaded across the gel. Integrated density of the western blot bands was normalized to actin and quantified using ImageJ software (National Institute of Health).

### Aβ_42_ treatment

Solution of oligomeric Aβ1–42 from lyophilized, high performance liquid chromatography-purified Aβ1– 42 was prepared as described previously. (Barghorn et al. 2005) First, 100% HFIP (1,1,1,3,3,3-hexafluoro-2-propanol) was used to reconstitute Aβ1–42 (1 mM), and then HFIP was removed by evaporation using a Speed Vac (Eppendorf, Hamburg, Germany). The obtained pellet was then resuspended to 5 μM in anhydrous dimethyl sulfoxide. This stock was diluted with phosphate-buffered saline to a final concentration of 400 μM, and sodium dodecyl sulfate was added to a final concentration of 0.2%. The resulting solution was then incubated at 37 °C for 18–24 hours. The preparation was again incubated at 37 °C for 18–24 hours after further dilution with phosphate-buffered saline to a final concentration of 100 μM. The nature of the Aβ1–42 oligomers of the preparation was then checked by sodium dodecyl sulfate–polyacrylamide gel electrophoresis. (Akhter et al. 2015) Cells were treated with 1 – 20 µM Aβ_42_ for 2 – 72 hours (data not shown). 5 µM Aβ_42_ treatment for 24 hours was selected for phagocytosis, apoptosis and inflammatory panel analyses because of moderate effect on cell survival and phagocytosis parameters (Fig 3C; Fig 4A).

### Morphology and Phagocytosis Assay

HMC3 morphology was examined under the microscope (20x Leica DFC3000G) with and without 5 µM Aβ_42_ treatment for 24 hours to visualize activated morphology (Fig 3A). Vybrant™ Phagocytosis Assay Kit (Invitrogen) was utilized to evaluate phagocytosis alterations in HMC3 cells with and without Aβ_42_ treatment and TREM2 overexpression or silencing. Cells were harvested from tissue culture plate and transferred to 96 well microplate at a concentration of 10^6^/ml. A Negative control was included consisting of only 200 µl of cell culture medium. A Positive control was included consisting of 200 ul of adjusted cell suspension. Also 200 ul of adjusted cell suspension was added to the experimental wells. The plates were incubated overnight optimum adherence. The next day cells were treated with 5μM Aβ_42_. After 0, 8 24, 48 and 72 hour incubation, media was removed and 100 µl was added of prepared fluorescent bioparticles to all the negative control, positive control and experimental wells and incubated for 2 hours. Bioparticles were removed and 100 µl of the prepared trypan blue suspension was added to all of the wells and incubated for 1 min at room temp. After removal of the trypan blue suspension phagocytosis status was immediately evaluated by fluorescence plate reader at 480 nm excitation, 520 nm emission (Synergy HTX multi-mode reader by BioTeK) (Fig 3C-D). (Gopallawa et al. 2020)

### Cell Survival and Apoptosis Assays

HMC3 cell viability in response to Aβ42 treatment was determined using MTT (0.5 mg/ml) assay and trypan blue exclusion (TBE). HMC3 cells were cultivated at a density of 10,000 cells/well in 96-well plates for 0, 8 24, 48 and 72 hours. Cells were then stained with 20 µl MTT 5 mg/ml stock solution (Thiazolyl Blue Tetrazolium Bromide: Sigma-Aldrich Cat: 5655) for 4 hours and then dissolved in DMSO. The optical density was determined at 590 nm (Plate Reader Synerty MX by BioTek). To evaluate whether the intrinsic apoptotic pathway is activated by 24 hours of 5 µM Aβ_42_ treatment with TREM2 overexpression or silencing, the MILLIPLEX® MAP 6-plex Bcl-2 Family Apoptosis Panel 1 Magnetic Bead Kit (Millipore: Cat. No. 48-682MAG) was utilized. Six analytes were measured (Luminex): pBAD (Ser112), BAD (total), BIM (total), BAX (total), Bcl-xL/BAD, and Mcl-1/BIM according to manufacturer instructions. This panel was selected because these Bcl-2 family members together provide a marker of the activation of a ‘life/death switch’ in cells. (Tsujimoto and Shimizu 2000)

### Inflammatory Factor Assay

Inflammatory factors were measured utilizing a MILLIPLEX MAP multiplex kit (HCYTMAG60PMX41BK; EMD Millipore) following the manufacturer’s instructions for human cytokine/chemokine analyte detection and the Luminex LX-200 system. The 38 inflammatory markers in the panel were as follows: epidermal growth factor (EGF), fibroblast growth factor 2 (FGF-2), eotaxin, TGF-α, G-CSF, FMS-like tyrosine kinase 3 ligand (Flt-3L), GM-CSF, fractalkine (also known as CX3CL1), IFN-α2, IFN-γ, growth-regulated oncogene (GRO), IL-10, monocyte chemotactic protein-3 (MCP-3, also known as CCL7), IL-12 40 kDa (IL-12p40), macrophage-derived chemokine (MDC), IL-12 70 kDa (IL-12P70), IL-13, IL-15, soluble CD40-ligand (sCD40L), IL-17A, IL-1 receptor agonist (IL-1RA), IL-1α, IL-9, IL-1β, IL-2, IL-3, IL-4, IL-5, IL-6, IL-7, IL-8, IFN-γ inducible protein 10 kDa (IP-10), MCP-1 (also known as CCL2), macrophage inflammatory protein (MIP) 1α (MIP-1α, also known as CCL3), MIP-1β (also known as CCL4), TNF-α, TNF-β, and vascular endothelial growth factor (VEGF).

## 3. Results

### HMC3 cells express moderately low TREM2 levels relative to other cell lines

For selection of human cell line with low to moderate TREM2 expression, TREM2 RNA and protein expression was evaluated in eight human untreated cell lines: HepG2 hepatocytes; neuronal IMR-32 and CHP212; glial U87, U118, HMC3; monocytes, THP1, U937. TREM2 relative to ACTB RNA expression was lower in HMC3 than most other cell lines (Fig 1A). TREM2 relative to Actin B was lower than many of the other cell lines (Fig 1B). TREM2 protein relative to RNA was moderately low compared to the other cell lines (Fig 1C).

**Fig 1.**
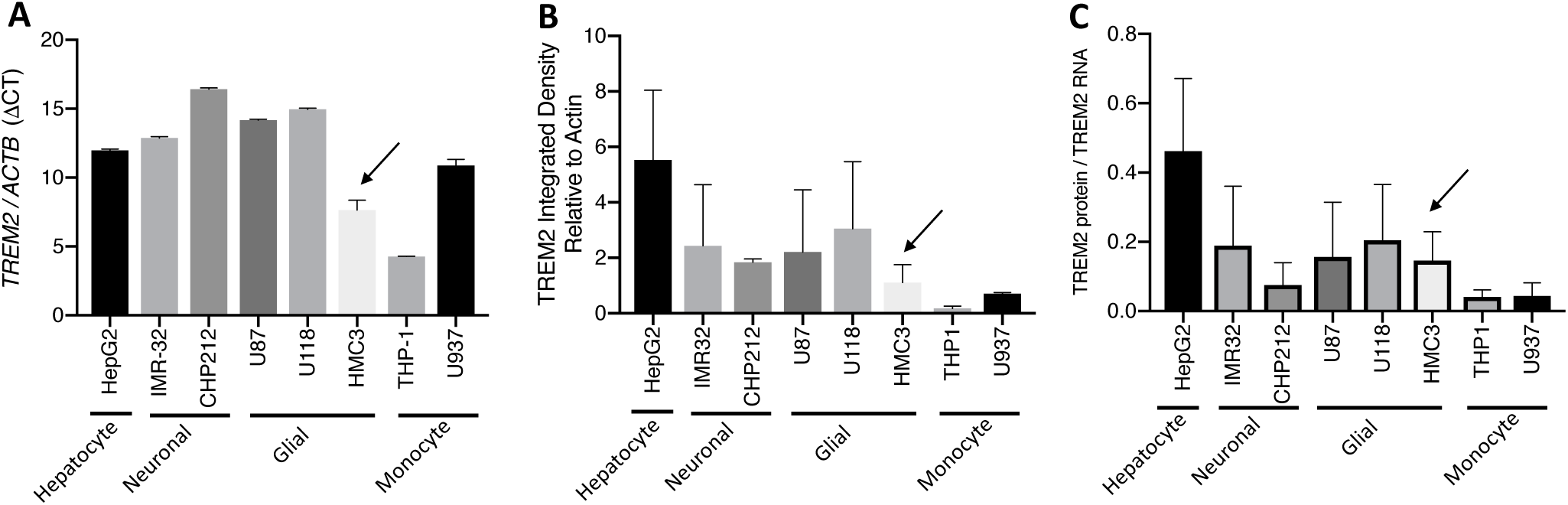
HMC3 cells express low to moderate TREM2 levels relative to other cell lines. *TREM2* RNA ΔCT levels (Relative to *ACTB*) are lower than most other cell lines (A). TREM2 protein (Relative to Actin B) are lower than most other cell lines (B). TREM2 protein relative to TREM2 RNA is lower the HepG2 cell line (C).

### Aβ_42_ treatment of HMC3 cells increases TREM2

To determine if Aβ_42_ treatment influences TREM2 expression in HMC3 cells, cells were treated and evaluated using immunostaining and Western blot. DAPI (blue), IBA1 (green) and TREM2 (red) immunostain of HMC3 cells treated with 5 μM Aβ_42_ indicates an increase in membrane cell surface TREM2 after 24 hours (Fig 2A). TREM2 full length 28 kDa is significantly elevated after 24 hours 5 μM Aβ_42_ treatment on Western blot (Fig 2B). TREM2 overexpression in HMC3 significantly increases TREM2 protein levels with or without Aβ_42_ treatment while TREM2 siRNA knockdown depletes TREM2 with or without Aβ_42_ treatment (Fig 2C). These results demonstrate that the Aβ_42_ peptide stimulates TREM2 protein expression in HMC3 cells.

**Fig 2.**
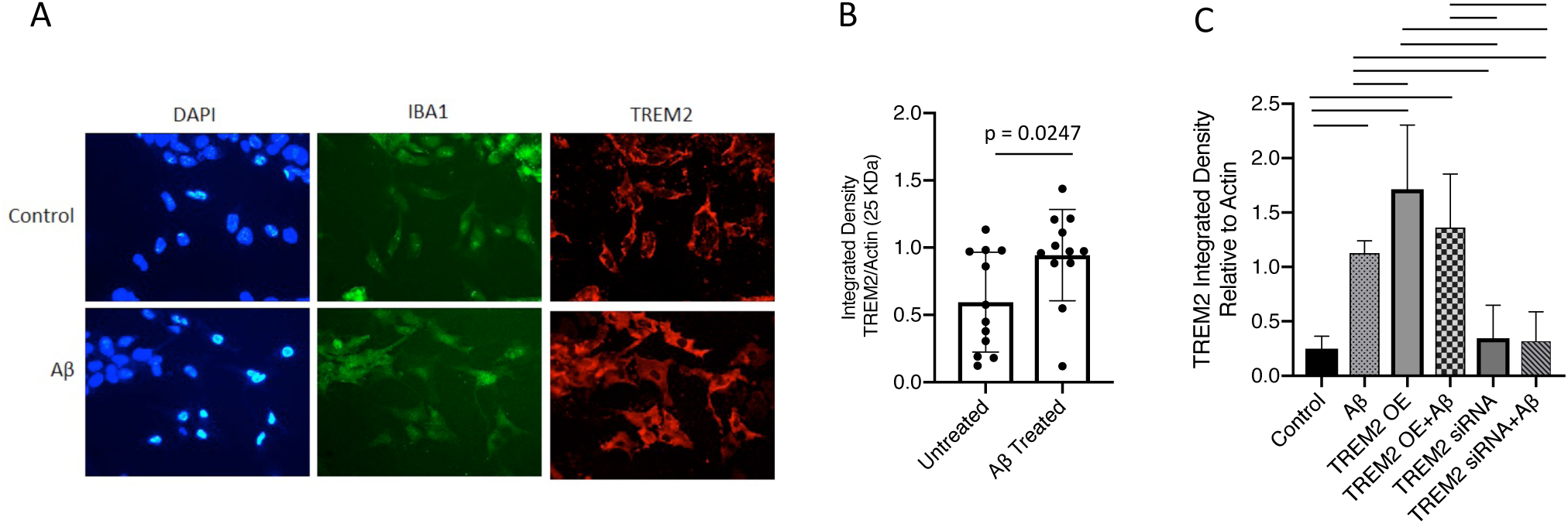
Aβ_42_ treatment of HMC3 cells increases TREM2. Treatment of HMC3 cells with 5 μM Aβ for 24 hours increases cell surface TREM2 expression (A). TREM2 full length protein expression is significantly increase after 24 hour 5 μM Aβ_42_ treatment (25 kDa: p-value = 0.0247) (B). In a separate experiment, full length TREM2 relative to Actin (Western Blot: 25 kDa) is significantly increased, upon 24 hour 5 μM Aβ_42_ treatment while overexpression of TREM2 (OE) plus Aβ_42_ treatment does not significantly increase TREM2. Knockdown of TREM2 (siRNA) significantly depletes TREM2 with and without 24 hour 5 μM Aβ_42_ treatment **(C)**. Fisher’s LSD test without correction for multiple comparisons significant p-value <0.05.

### Lack of TREM2 decreases Aβ_42_ stimulated phagocytosis in HMC3 cells

To determine if TREM2 influences HMC3 Aβ_42_ stimulated phagocytosis, HMC3 cells were treated with 5 μM Aβ_42_ for 24 hours and phagocytosis assays performed. Aβ_42_ treatment changed HMC3 cell morphology from ramified to amoeboid after 24 hours (Fig 3A-B). Aβ_42_ treatment significantly increased % phagocytosis at 8-24 hours and declined after 48-72 hours (Fig 3C). TREM2 overexpression significantly increases phagocytosis but does not enhance phagocytosis after 24 hours of 5 μM Aβ_42_ treatment while TREM2 knockdown decreases phagocytosis with or without 24 hour 5 μM Aβ42 treatment (Fig 3D). These results demonstrate that a lack of TREM2 depletes Aβ_42_ stimulated phagocytosis in HMC3 cells.

**Fig 3.**
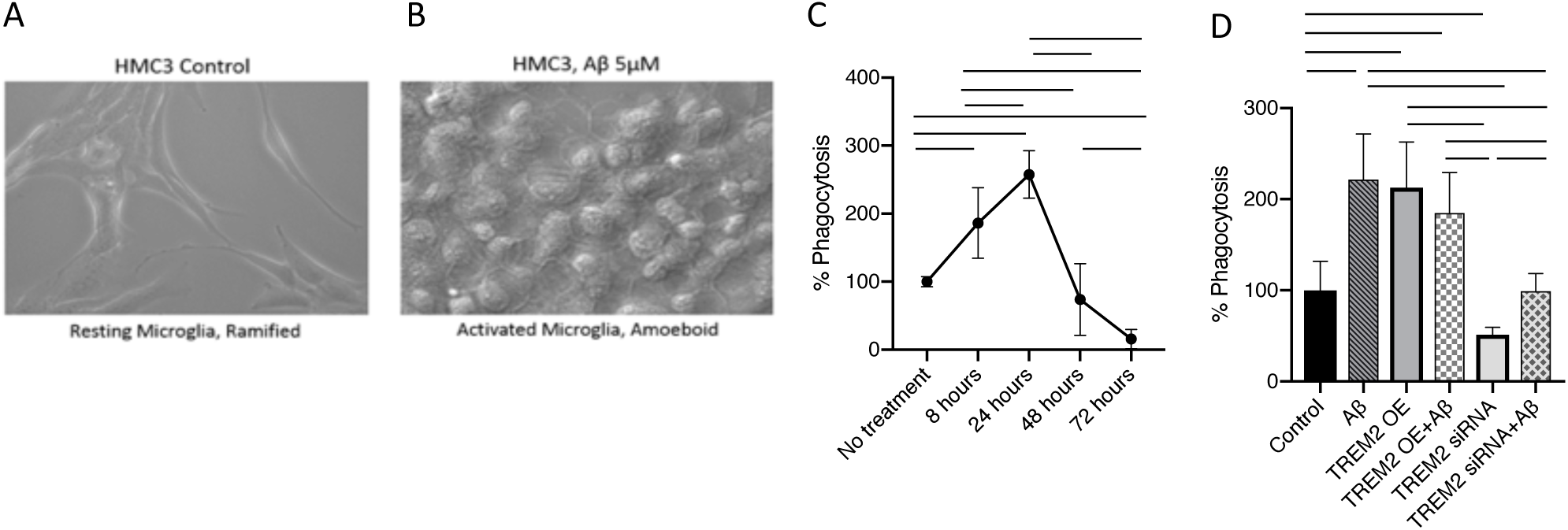
Lack of TREM2 decreases Aβ_42_ stimulated phagocytosis in HMC3 cells. Treatment of HMC3 cells with 5 μM Aβ_42_ for 24 hours changes cell morphology from resting ramified microglial **(A)** to activated amoeboid microglia **(B).** Maximum phagocytosis is observed after 24 hours of 5 μM Aβ_42_ treatment (lines represent significant p-values <0.01) **(C).** After 24 hours of 5 μM Aβ_42_ treatment there is a significant increase in phagocytosis with or without overexpression of TREM2 and upon TREM2 knockdown there is a decrease in phagocytosis with or without 24 hour 5 μM Aβ_42_ treatment **(D).**

### Aβ_42_ treatment alters cell viability in HMC3 cells

To understand the influence of Aβ_42_ treatment on HMC3 cell survival over time. Cells were treated with 5 μM Aβ_42_ for 72 hours and an MTT assay for cell viability was performed at 8, 24, 48 and 72 hours. After 8 hours there is a significant decline in viability (p-value <0.001) suggesting that both cell death and functional phagocytosis occurs at 24 hours of 5 μM Aβ_42_ treatment in HMC3 cells (Fig 4).

**Fig 4.**
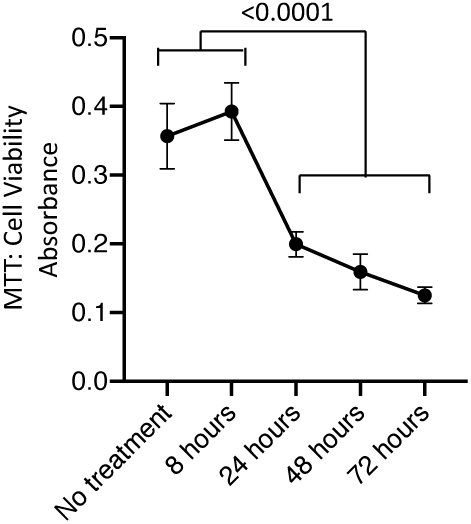
Aβ_42_ treatment alters cell survival, pro-apoptotic and anti-apoptotic factors in HMC3 cells. HMC3 cell viability significantly declines after 8 hours of 5 μM Aβ_42_ treatment Bars represent Fisher’s LSD test without correction for multiple comparisons significant p-value <0.05.

### TREM2 modulates apoptotic machinery in HMC3 cells exposed to the Aβ_42_ peptide

To determine if TREM2 plays a role intrinsic apoptotic response to 24 hours of 5 μM Aβ_42_ in HMC3 cells, the apoptosis-related proteins BAX, BAD, Bcl-xL/BAD, BIM, pBAD and MCL-1/BIM were evaluated. At 24 hours of Aβ_42_ treatment Bax, Bad, and pBad are reduced (Fig 5A-B). TREM2 overexpression (TREM2 OE) alone does not influence these apoptotic proteins while TREM2 silencing (TREM2 siRNA) alone increases Bax. (4) When TREM2 is overexpressed, 24 hour of Aβ_42_ treatment results in a significant increase in Mcl-1/Bim. When TREM2 is silenced, Bax, Bcl-xL/Bad are decreased in response to 24 hour Aβ_42_ treatment. In addition, at 24 hour Aβ_42_ treatment TREM2 silencing results in a decrease in Bax compared to TREM2 overexpression. These results suggest that Bax related apoptosis is influenced by TREM2-related activation of the Mcl-1/Bim complex.

**Fig 5.**
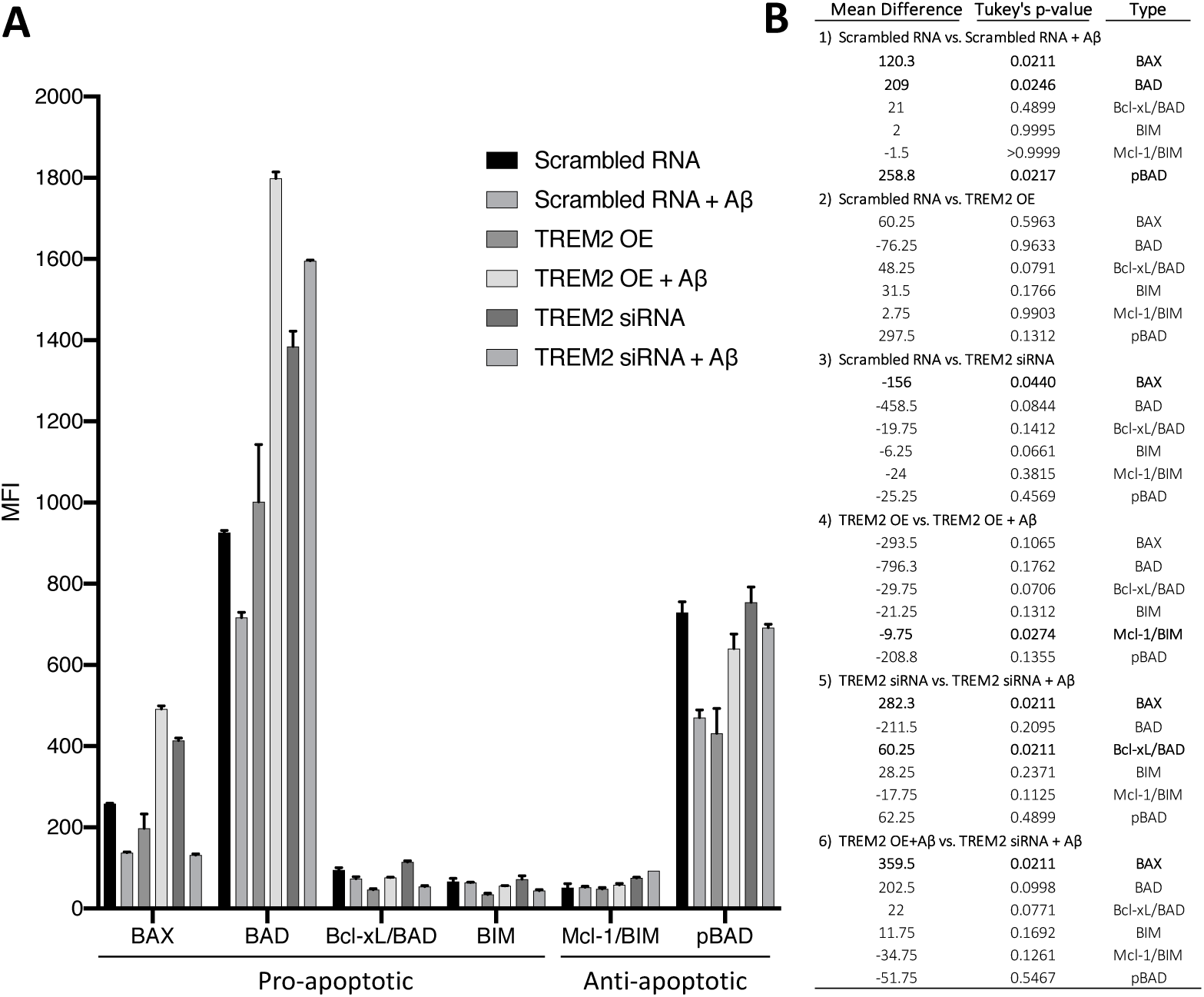
TREM2 modulates apoptotic machinery in HMC3 cells. **(1)** Bax, Bad, and pBad are reduced in response to 24 hour Aβ_42_ treatment. **(2)** TREM2 overexpression (TREM2 OE) does not influence these apoptotic proteins. **(3)** TREM2 silencing (TREM2 siRNA) increases Bax. **(4)** Mcl-1/Bim is increased in response to 24 hour Aβ_42_ treatment in the context of TREM2 overexpression. **(5)** Bax, Bcl-xL/Bad are decreased in response to 24 hour Aβ_42_ treatment in the context of TREM2 silencing. **(6)** In response to 24 hour Aβ_42_ treatment TREM2 silencing results in a decrease in Bax compared to TREM2 overexpression. **(A)** Significance (p-values) are Tukey adjusted for multiple comparisons and mean difference (Mean Diff.) is shown for each apoptotic factor. **(B)**

### Proinflammatory factors generally increased upon TREM2 overexpression in HMC3 cells

To determine if TREM2 plays a role in HMC3 inflammatory response to Aβ_42_ HMC3 cell media was collected and a panel of 38 cytokines and chemokines were evaluated after 24 hours of 5 μM Aβ_42_ treatment both with and without TREM2. Generally, anti-inflammatory and immunomodulatory factors (on the inflammatory panel) are below or near the lower limit of detection in HMC3 cells. Only 6 inflammatory factors; FGF-2, IP-10, VEGF, MIP-1a, GRO and IL-8, are significantly associated with TREM2 mediated response to 24 hours of 5 μM Aβ_42_ treatment in HMC3 cells (Fig 6A-B). There is no change in inflammatory factors after 24 hour Aβ_42_ treatment alone. In addition, there is not a significant change in inflammatory factor level with TREM2 overexpression or silencing. Furthermore, when TREM2 is overexpressed there is not a significant difference in inflammatory factor level after 24 hour Aβ_42_ treatment. In contrast, when TREM2 is silenced, FGF-2 is increased and MIP-1a is decreased after 24 hour Aβ_42_ treatment. In addition, TREM2 silencing results in a decrease in IP-10, MIP-1a, IL-8 and an increase in FGF-2, VEGF, GRO, compared to TREM2 overexpression, in response to Aβ_42_ treatment. This suggests that in HMC3 cells TREM2 plays a role in Aβ induced activation of IP-10, MIP-1a, and IL-8 as well as the inhibition of the FGF-2, VEGF, GRO.

**Fig 6.**
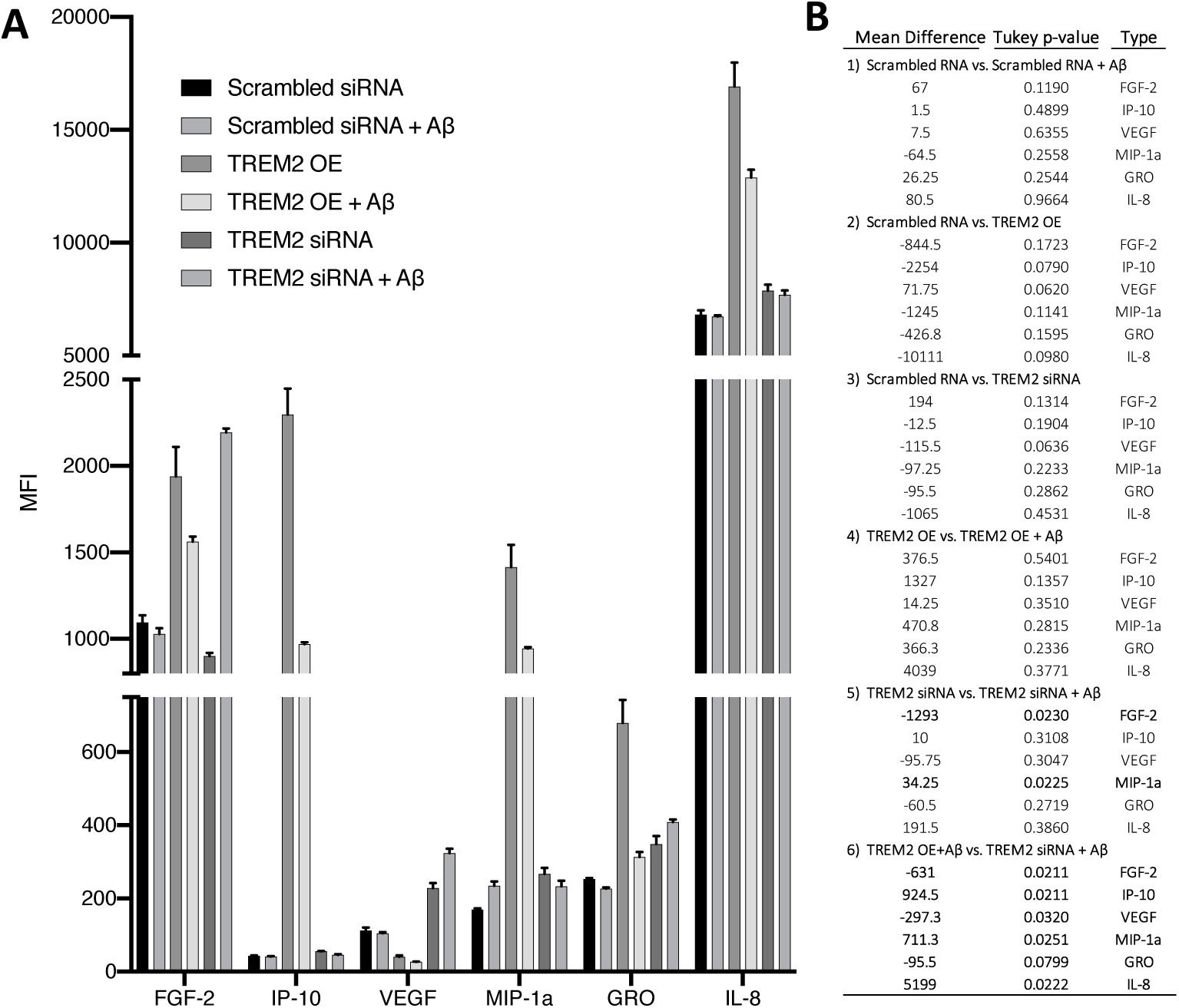
Proinflammatory factors generally increased upon TREM2 overexpression in HMC3 cells. **(1)** No change in inflammatory factors after 24 hour Aβ_42_ treatment. **(2)** No change in inflammatory factors with TREM2 overexpression. **(3)** No change in inflammatory factors when TREM2 silenced. **(4)** No change in inflammatory factors with TREM2 overexpression after 24 hour Aβ_42_ treatment. **(5)** FGF-2 is increased and MIP-1a is decreased when TREM2 silenced after 24 hour Aβ_42_ treatment. **(6)** In response to 24 hour Aβ_42_ treatment TREM2 silencing results in a decrease in IP-10, MIP-1a, IL-8 and a increase in FGF-2, VEGF, GRO, compared to TREM2 overexpression. **(A)** Significance (p-values) are Tukey adjusted for multiple comparisons and mean difference (Mean Diff.) is shown for each apoptotic factor. **(B)**

## 4. Discussion

TREM2 has been strongly associated with the microglia response to amyloid in mouse models of AD. Increasing evidence supports the role of TREM2 in microglial function where phagocytosis, apoptosis and the inflammatory response have all been linked to TREM2. However, missing information about which specific apoptotic and inflammatory proteins are influenced by TREM2 in response to Aβ_42_ is less well characterized in human cell lines. To test the hypothesis that TREM2 influences the response to Aβ toxicity by enhancing phagocytosis and inhibiting both the BCL-2 family of apoptotic proteins and pro-inflammatory cytokines, the human HMC3 microglial cell line was selected because it has a moderate expression level of endogenous TREM2 (Fig 1). This is necessary for experiments that involve both overexpression and silencing of TREM2.

Given previous evidence that there is an increase in TREM2 with Aβ_42_ treatment in mouse microglia or in AD mouse models (Brendel et al. 2017, Ulrich et al. 2017, Zhao et al. 2018), but TREM2 has not been characterized in HMC3 cells, we first addressed the question of the whether Aβ_42_ treatment increased TREM2 in this human microglia cell line. TREM2 immunohistochemistry indicates that TREM2 is elevated upon Aβ_42_ treatment after 24 hours (Fig 2A). This increase in TREM2 was also observed by western blot derived protein expression in (Fig 2B). When TREM2 is overexpressed in HMC3 cells, this overexpression is maintained upon 24 hour Aβ_42_ treatment and depleted upon TREM2 silencing (Fig 2C). This confirms that we can effectively overexpress and silence TREM2 in this cell line under conditions of Aβ_42_ toxicity as previously demonstrated in mouse cell lines (Jiang et al. 2014, Zhao et al. 2018).

To demonstrate that HMC3 cells undergo alterations in morphology upon exposure to Aβ_42_ treatment as previously described in other cell lines (Pan et al. 2011, Hansen et al. 2018), they were treated for 24 hours with 5 µM Aβ_42_. The cells normally have a resting ramified phenotype (Fig 3A) and then when stimulated by 24 hours of 5 µM Aβ_42_ they develop an activated amoeboid phenotype (Fig 3B). TREM2 regulates microglial cholesterol metabolism upon chronic phagocytic challenge (Nugent et al. 2020) and recognizes anionic ligands on bacteria and some eukaryotic cells. (Daws et al. 2003) This amoeboid phenotype in response to Aβ has been described in mouse cell lines (Krabbe et al. 2013, Caldeira et al. 2017) but has not been described previously in HMC3 cell lines. To quantify the change phagocytic activity over time, a phagocytosis assay using the fluorescein labeled bioparticles was performed (Fig 3C). A significant increase in phagocytosis in response to 5 µM Aβ_42_ treatment occurs until 24 hours and then significantly declines. This suggests that in HMC3 cells increase phagocytic response to Aβ for 24 hours. However, there is an eventual decline in this phagocytic response to Aβ in this cell line with low to moderate endogenous TREM2 (Fig 3C). However, at 24 hours, when TREM2 is overexpressed, there is an increase in phagocytosis compared to the control for both with and without Aβ42 treatment (Fig 3D). In contrast, when TREM2 is silenced, phagocytosis is inhibited and only reaches the level of the control upon 24 hour Aβ_42_ treatment (Fig 3D). Together, this suggests that TREM2 is involved in phagocytosis. But, given the observed low level of phagocytosis upon TREM2 silencing in conjunction with Aβ_42_ treatment, it is possible there is another active phagocytosis mechanism in this cell line. This is not novel information in the context of myeloid cell TREM2 (Hsieh et al. 2009, Kawabori et al. 2015) and other Aβ-related non-TREM2 phagocytic mechanisms have been described in myeloid cells (Lai and McLaurin 2012, Yamanaka et al. 2012). However, this is new information about HMC3 cells. TREM2 mutations disrupt phagocytic capabilities of microglia and other cell types. (Kleinberger et al. 2014, Kawabori et al. 2015, Schlepckow et al. 2017, Yao et al. 2019) Results of this study indicate that without TREM2 there is a significantly reduced phagocytic response to Aβ.

The Bcl-2 family of apoptotic proteins have been implicated in the Bcl-2 family Akt signaling response to amyloid. (Clementi et al. 2006, Yin et al. 2011, Shang et al. 2012, Zhu et al. 2015) However, even though TREM2- related Akt signaling has been described (He et al. 2019, Chen et al. 2020) to our knowledge a link between TREM2 specific signaling and Bcl-2 apoptosis has not been studied. Aβ_42_ treatment influences cell survival in HMC3 cells where after 8 hours there is a significant decline in viability (Fig 4). Interestingly, at 24 hours there is an increase in phagocytosis suggesting that even though there is a decline in viability at 24 hours there is still functional phagocytosis. Taken together this suggests that cell death occurs between 8 and 24 hours but the cells are still functional in that timeframe where they are phagocytic until somewhere between 24 and 48 hours. Therefore, next to understand if the intrinsic apoptotic machinery is influenced by 24 hours of Aβ_42_ treatment, the Bcl-2 family of apoptotic factors were evaluated in HMC3 cells. Bax, Bad, and pBad are reduced in response to 24 hours of Aβ_42_ treatment. TREM2 overexpression alone does not influence the Bcl-2 family of apoptotic proteins in HMC3 cells. Interestingly, Mcl-1/Bim is increased in response to 24 hour Aβ_42_ treatment in the context of TREM2 overexpression suggesting reduced apoptosis due to sequestration of pro-apoptotic Bim by antiapoptotic Mcl-1. In contrast, TREM2 silencing increases Bax while Bad and Bcl-xL/Bad are decreased in response to 24 hour Aβ_42_ treatment in the context of TREM2 silencing. Furthermore, in response to 24 hour Aβ_42_ treatment TREM2 silencing results in a decrease in Bax compared to TREM2 overexpression. Taken together, this novel information suggests that TREM2 inhibits Aβ induced apoptosis by activating the Mcl-1/Bim complex and thus inhibiting the production of Bax in HMC3 cells.

TREM2 inhibits the inflammatory response to LPS in mouse microglia by suppressing the PI3K/NF-κB signaling, (Li et al. 2019) but less is known about the response to Aβ in human microgliaTREM2 rare genetic variants located in the phagocytic receptor triple the risk of developing AD (Jonsson et al. 2013, Abduljaleel et al. 2014) highlighting the importance of TREM2 in brain homeostasis. Critical cellular biology studies link TREM2 downstream signaling to inflammatory factors. (Li et al. 2019, Long et al. 2019, Wang et al. 2019, Zhu et al. 2020) However, which inflammatory factors are influenced by TREM2 after Aβ treatment had not been fully elucidated. Since a main function of microglia is to secret inflammatory factors and the inflammatory factors influenced by TREM2 and Aβ_42_ treatment have not been characterized in HMC3 cells, we analyzed a panel of 38 inflammatory factors. Interestingly, there is not a significant change in inflammatory factors after 24 hour Aβ_42_ treatment. In addition, there is not a significant change in inflammatory factors when TREM2 is overexpressed or silenced. Furthermore, there is no change in inflammatory factors with TREM2 overexpression after 24 hour Aβ_42_ treatment. However, FGF-2 is significantly increased and MIP-1a is decreased after 24 hour Aβ_42_ treatment when TREM2 is silenced. In response to 24 hour Aβ_42_ treatment TREM2 silencing results in a decrease in IP-10, MIP-1a, IL-8 and an increase in FGF-2, VEGF, GRO, compared to TREM2 overexpression. These results suggest that under conditions of Aβ toxicity TREM2 is involved in activation of IP-10, MIP-1a and IL-8 and the inhibition of FGF-2, VEGF, and GRO, in HMC3 cells. Interestingly, IP-10, MIP-1a and IL-8 all play a role in chemoattraction and cell recruitment (Engelhardt et al. 1998, Turner et al. 2014)while FGF-2, VEGF, and GRO play a role in response to injury, angiogenesis, wound healing, and the immune response (Werner and Grose 2003, Frantz et al. 2005) suggesting that TREM2 is involved in activating chemoattraction and recruitment based cell biology, while inhibiting the response to injury.

In conclusion, this study has elucidated the relationship between TREM2 and HMC3 microglial biology in response to neurotoxic Aβ. A robust phagocytic response to Aβ requires TREM2 in these cells. TREM2 inhibits Aβ induced apoptosis by activating the Mcl-1/Bim complex and thus inhibiting the production of Bax in HMC3 cells. Furthermore, in response to Aβ toxicity, TREM2 is involved in activation of IP-10, MIP-1a and IL-8 which play a role in chemoattraction and cell recruitment. In contrast, TREM2 is involved in inhibition of the FGF-2, VEGF, and GRO which play a role in response to injury, angiogenesis, wound healing, and the immune response. This study reveals novel insights into contribution of TREM2 in the response to Aβ in microglia and suggests that therapeutics that seek to target TREM2 biology may have a broad influence on microglial function that warrants further investigation.

## Authorship

All authors contributed to the design of the experiments, analysis of the data, data interpretation and writing the manuscript.

## Declaration of Competing Interest

The authors declare no conflict of interest.

## Acknowledgements

This study was supported by the National Institute of Health R00AG034214 and R56 AG063870.

## Notes

### Competing Interest Statement

The authors have declared no competing interest.

